# Automatic mutual information noise omission (AMINO): generating order parameters for molecular systems

**DOI:** 10.1101/745968

**Authors:** Pavan Ravindra, Zachary Smith, Pratyush Tiwary

**Affiliations:** Department of Chemistry and Biochemistry, University of Maryland, College Park 20742, USA; Department of Computer Science, University of Maryland, College Park 20742, USA; Biophysics Program, University of Maryland, College Park 20742, USA; Institute for Physical Science and Technology, University of Maryland, College Park 20742, USA

## Abstract

Molecular dynamics (MD) simulations generate valuable all-atom resolution trajectories of complex systems, but analyzing this high-dimensional data as well as reaching practical timescales even with powerful super-computers remain open problems. As such, many specialized sampling and reaction coordinate construction methods exist that alleviate these problems. However, these methods typically don’t work directly on all atomic coordinates, and still require previous knowledge of the important distinguishing features of the system, known as order parameters (OPs). Here we present AMINO, an automated method that generates such OPs by screening through a very large dictionary of OPs, such as all heavy atom contacts in a biomolecule. AMINO uses ideas from information theory and rate distortion theory. The OPs learnt from AMINO can then serve as an input for designing a reaction coordinate which can then be used in many enhanced sampling methods. Here we outline its key theoretical underpinnings, and apply it to systems of increasing complexity. Our applications include a problem of tremendous pharmaceutical and engineering relevance, namely, calculating the binding affinity of a protein-ligand system when all that is known is the structure of the bound system. Our calculations are performed in a human-free fashion, obtaining very accurate results compared to long unbiased MD simulations on the Anton supercomputer, but in orders of magnitude less computer time. We thus expect AMINO to be useful for the calculation of thermodynamics and kinetics in the study of diverse molecular systems.

## I. INTRODUCTION

Molecular Dynamics (MD) has become a routine tool for simulating and understanding the various structural, thermodynamic and kinetic properties of complex real-world molecular systems. MD simulations make it possible to study these systems comprising millions of atoms reaching timescales of microseconds and beyond, while maintaining all-atom spatial and femtosecond temporal resolutions. In spite of this staggering success of MD over the decades, one is faced with two central and inter-connected problems. First, these long simulations with the aforementioned high spatial and temporal resolution can easily produce an overwhelming amount of data (easily into terabytes). How do we make sense of this data? Second, even these long simulations are not long enough, as currently in spite of the best available supercomputing resources MD can reach at best a microsecond in practical wall-clock time. Thus specialized so-called “enhanced sampling” algorithms^1^ have been developed that allow us to reach much longer timescales of seconds, minutes and beyond in a statistically accurate manner.

While these two challenges to MD might appear disconnected, they share a commonality – namely, the need for dimensionality reduction. The complex, often bewildering dynamics that seems to happen in an extremely high-dimensional configuration space, can often be reduced to certain key variables that capture the underlying physics or chemistry. The other remaining variables are then either irrelevant or can simply be mapped into noise. These key variables, which we refer to as “order parameters (OP)” can then serve to form a data-efficient low-dimensional picture of complicated molecular systems and processes. These OPs serve as internal coordinates that are useful as a basis set for the description of the processes of interest.^2^ Through one of many available methods^3–7^, they can then also be mixed into an even lower-dimensional reaction coordinate (RC). By then enhancing fluctuations along this RC, enhanced sampling methods such as metadynamics^8^ and umbrella sampling^9^ can tackle the second challenge mentioned above, i.e. assessing processes that happen far slower than the capabilities of unbiased MD. From the above discussion it should thus be clear that the proper selection of OPs is crucial for analysis and enhancing of MD simulations.^5,8^

In this work we propose a new fairly automated computational scheme for the selection of such OPs. Previously, selection of OPs has relied purely on biophysical intuition or knowledge of the system and processes of interest.^5^ However, for novel systems of interest, there may not be enough information about the system to make claims about which OPs are relevant to a particular chemical process. Creating a robust, flexible algorithm to screen for OP redundancy is thus a problem of great interest to the field of MD simulations. Selecting noisy OPs that are not pertinent to the process of interest or OPs that provide redundant information can slow down calculations or even yield misleading results. For instance consider the case where the selected OPs are used to construct a RC for metadynamics simulation (or other biased simulations).^8^ Providing redundant, correlated or noisy OPs can lead to an inefficient biasing protocol that might end up being even slower than unbiased MD. While the developed formalism and algorithms should be quite generally applicable to a variety of real-world molecular systems, here we consider as an illustrative test-case the calculation of absolute binding affinity of a protein-ligand system in explicit water.

Our algorithm, which we name “Automatic Mutual Information Noise Omission (AMINO)”, uses a mutual information based distance metric to find a set of minimally redundant OPs from a much larger set, and then uses K-means clustering with this distance metric, together with ideas from rate distortion theory^10^ to find representative OPs from each cluster that provide maximum information about the given system. We demonstrate the effectiveness of our method on analytical model systems and the much larger FKBP-BUT protein-ligand system. In each example we begin with absolutely no prior information on the system other than an initial set of coordinates for each atom. We apply AMINO to generate a set of OPs from an unbiased trajectory of the system, then we generate a reaction coordinate using SGOOP to run metadynamics, enhancing the dissociation process and accurately calculating the absolute binding free energy.

We believe that the current work is arguably one of the first illustrations of a fully automated pipeline where starting with a known protein data bank (PDB) structure of a bound protein-ligand system, a force-field, and a short MD run exploring the bound pose (but not necessarily showing dissociation), one performs enhanced sampling of the dissociation and obtains binding free energy in order of magnitude speed-up relative to unbiased MD. The OPs generated by AMINO can be used in any other procedure of choice for generating reaction coordinate, such as TICA,^3^ RAVE,^6^ or VAC^7^, followed by use in an enhanced sampling protocol not limited to metadynamics. The OPs identified by AMINO also form a most concise dimensionality reduction of an otherwise gargantuan MD trajectory. Selecting OPs has been a major barrier toward creating a fully automated enhanced sampling procedure, and now, AMINO can serve as an automated protocol for selecting OPs on an information theoretic basis. We thus expect AMINO to be useful to a wide range of practitioners of molecular simulations.

## II. THEORY

We start this section by summarizing the key steps in our algorithm, and then gradually go through every step in detail. Our ultimate goal is to construct clusters of similar OPs and choose a single OP from each cluster that best describes its parent cluster as a whole. The number of clusters or equivalently the number of OPs is learned on-the-fly through a formalism based on rate distortion theory. The key input to our algorithm is a large set of OPs and their time series in a short unbiased trajectory. Note that AMINO does not need temporal information on the OPs, as such the time series could be coming from independent unbiased simulations, appropriately reweighted biased simulations, or could even be temporally scrambled. Given this input, AMINO involves the following sequential procedure:

1. Cluster the input OPs using a mutual information variant that serves as a distance metric. The idea here is that not all OPs carry significantly different information, and with an appropriate distance function, we can identify groups of OPs carrying similar information.
2. Select a single OP from each cluster to describe best all of the OPs within the cluster. This OP is thus most representative of the information carried by all OPs in the cluster that it belongs to.
3. Finally, determine the appropriate total number of clusters, or equivalently the number of OPs to use to describe the entire set of OPs that AMINO started out with.

In step 1, we use a mutual information based distance function to measure the similarity of any two OPs.^11^ In step 2, we apply a variation on the well-known K-Means clustering algorithm,^12^ and finally in step 3, apply a recent implementation of rate distortion theory to clustering in order to find the ideal number of clusters or OPs.^13^

### A. A Distance Function based on Mutual Information

Any clustering procedure requires a relevant distance metric. In Cartesian coordinate systems, the most commonly used metric is the Euclidean distance between two points.^12^ However, there is no easily identifiable low-dimensional space that OPs lie upon in which the Euclidean distance between two OPs would shed meaningful information about the similarity of the OPs. Thus, we need to define a new metric where the distance between two OPs would correlate with how similar they are to each other in terms of information they carry.

More rigorously speaking, a useful distance metric for this setting satisfies all of the relevant properties of any other distance metric in addition to requirements specific to this system. Together these requirements are detailed below for a distance metric *D*(*X, Y*) for any given pair of OPs *X* and *Y*.

1. Non-negativity: *D*(*X, Y*) ≥ 0
2. Symmetry: *D*(*X, Y*) = *D*(*Y, X*)
3. OPs *X, Y* that are close to each other, as determined by small *D*(*X, Y*), should be “redundant”. This would mean that knowledge of one OP’s time series provides substantial information about the time series of the other OP. In the clustering scheme, these parameters should be clustered together so that only one of the two is likely to be chosen for the final set.
4. OPs *X, Y* that are far from each other, as determined by large *D*(*X, Y*), should be relatively independent. This would mean that they are not redundant and should not be clustered together.

The Mutual Information (MI) based distance metric proposed by Kraskov and co-workers satisfies all of the above requirements.^11^ This distance function is a variant of Mutual Information normalized over the range [0, 1].

In the continuous setting,

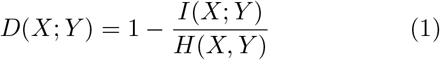

where *I* is the mutual information between *X* and *Y* and *H* is the joint entropy of *X* and *Y*, which are well-defined terms in information theory detailed for instance Ref. 10. For binned probability distributions over the OP’s time series, Eq. 1 can be expressed as:

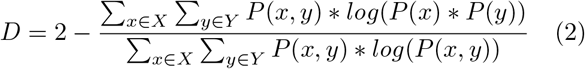

Eq. 1 and Eq. 2 are equivalent formulations of the MI-based distance between two OPs *X* and *Y*. Inspection of Eq. 2 gives an intuitive reasoning about the meaning of *D*. The denominator of the fraction component of Eq. 2 contains the joint probability distribution of *X* and *Y* while the numerator is the joint probability distribution assuming independence between *X* and *Y*. The core purpose of this equation is to compute how different the true join probability distribution is from the independence assumption.

Clustering using the mutual information based distance metric results in clusters of OPs with low mutual distances, where knowing the time series of a single OP in a cluster would give significant information about the trajectories of the other OPs in the cluster. Any two OPs from different clusters, however, would be fairly independent, and knowledge of either OP’s time series would provide little information about the other time series.

### B. Dissimilarity Matrix and Clustering

Now that we have defined a suitable distance metric in Sec. II A, we are ready to complete the first step in AMINO, namely clustering of different OPs by how similar or dissimilar they are. K-Means clustering provides a powerful approach to group a set of data points into *k* clusters for some provided number of clusters *k*. An overview pseudocode for K-Means clustering is provided in Algorithm 1. We stress that Algorithm 1 must be provided with the number of clusters *k* and that determining the optimal *k* for a given set of data is a separate (yet extremely) important concern. In Sec. II C, we discuss this problem in detail along with our solution to it.

As stated in Algorithm 1, the initial centroids for K-Means clustering are randomly generated points in the same space as the elements of S, the set of data points to be clustered. For traditional K-Means clustering, this initial randomization is acceptable due to the flexible nature of the centroids. For a proper selection of *k*, even if two of the randomly generated initial centroids are very close to each other, some other cluster in the data set that is unrepresented by a centroid will “pull” one of the centroids towards itself over time, as demonstrated in Fig. 1. Fig. 1 (a)–(c) show the time evolution for a set of data points using traditional K-Means clustering. In the first iteration (Fig. 1(a)), the centroids *µ*_1,…,*k*_ were chosen as a random set of *k* points in the same space as the points in the data set *S*. It is important to note that the centroids *µ*_1,…,*k*_ are not directly sampled from *S* itself, so there is no guarantee that these points will be near any of the clusters in *S*.

**FIG. 1:**
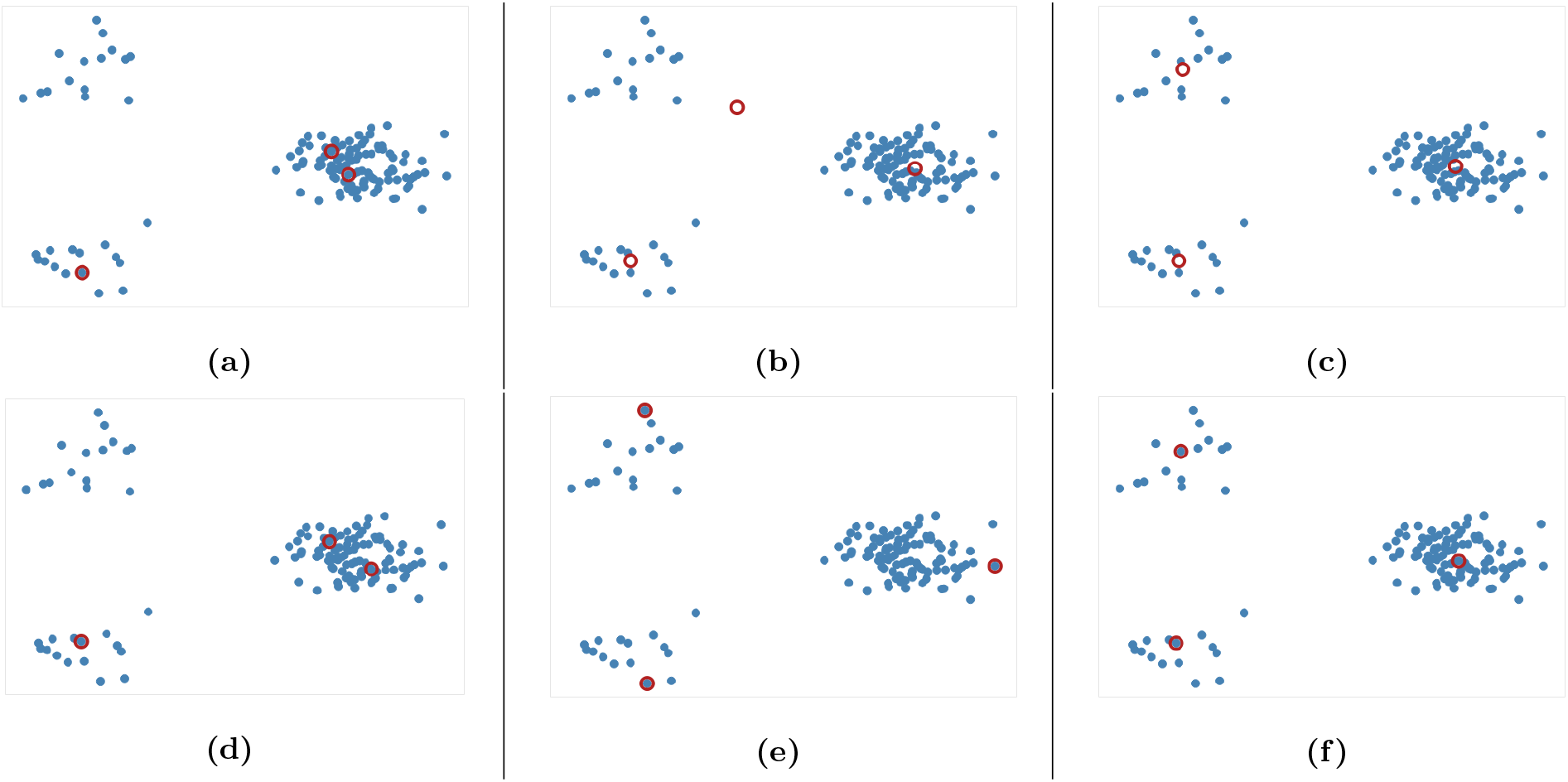
(a)-(c) Traditional K-Means clustering (Algorithm 1) over multiple iterations for a dataset. (d)-(f) Important figures for Restricted Centroid K-Means clustering (Algorithm 2) Red circles represent centroids, while blue points represent points in the dataset. (a) The initial random centroids. The two rightmost centroids are closest to the same cluster. (b) An intermediate state where one of the centroids is moving from one cluster to another. Note that the bottommost centroid is far from any of the points in the original data set. (c) The final set of centroids that the K-Means algorithm converges to. (d) An example initial distribution of centroids where the centroids would stay“stuck” in a cluster if a restricted centroid approach is used. The provided centroids are the final centroids using the restricted centroid K-Means algorithm, meaning that additional iterations will return the same set of centroids and simply not give 3 centroids in 3 clusters. (e) An initial distribution of centroids that have been chosen to maximize internal distance, or dissimilarity. (f) The result of running restricted centroid clustering (Algorithm 2) on the initial centroids from (e).

In the K-Means clustering example provided, two of the randomly chosen initial centroids are closest to the same cluster show in Fig. 1(a), yet by the second iteration (Fig. 1(b)), one of the centroids is beginning to move to the unrepresented cluster. In traditional K-Means clustering, the centroids do not need to take on values from the original data set *S*, so centroids are flexible and can move from one cluster to the other. This transition of a centroid from one cluster to another allows for traditional K-Means Clustering to converge to a reasonable solution, which is shown in Fig. 1(c).

However, now consider the problem in this work, where the points to be clustered represent OPs. Initial centroids cannot be generated randomly from the space of OPs, as all centroids must be points within the set of provided OPs. The same restriction applies to successive iterations-the updated centroid for a given cluster should be the element of the cluster that is closest to the remaining points. We thus need to revisit the definition of a centroid itself, and here we use the internal distortion of a cluster for a given candidate centroid to measure how well the candidate centroid represents the entire cluster, and by minimizing the distortion, we get the centroid of a given set of points. More formally, the centroid of some cluster *z* in the original set of data points can be defined as:

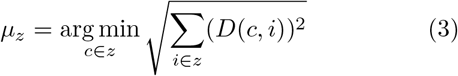

A pseudocode for K-Means clustering with restricted centroids is provided as Algorithm 2.

As a result of the requirement that the centroid must be an actual element of a cluster, we lose on the ability of K-Means where to transition a centroid from one cluster to another. In other words, centroids can become “stuck” within dense clusters of OPs. The resulting *k* clusters may fail to accurately represent clusters that are not captured by the initial set of randomly-generated starting centroids.

As an example, consider the initial centroids given in Fig. 1(d). These initial centroids are the final solution of restricted centroid K-Means clustering, meaning that K-Means algorithm has already converged to its solution. There is clearly a cluster that is unrepresented by any of the 3 centroids, yet due to the decreased centroid flexibility, neither of the centroids that share a cluster will cross the gap between the clusters in the data set.

#### Algorithm 1 K-Means Clustering

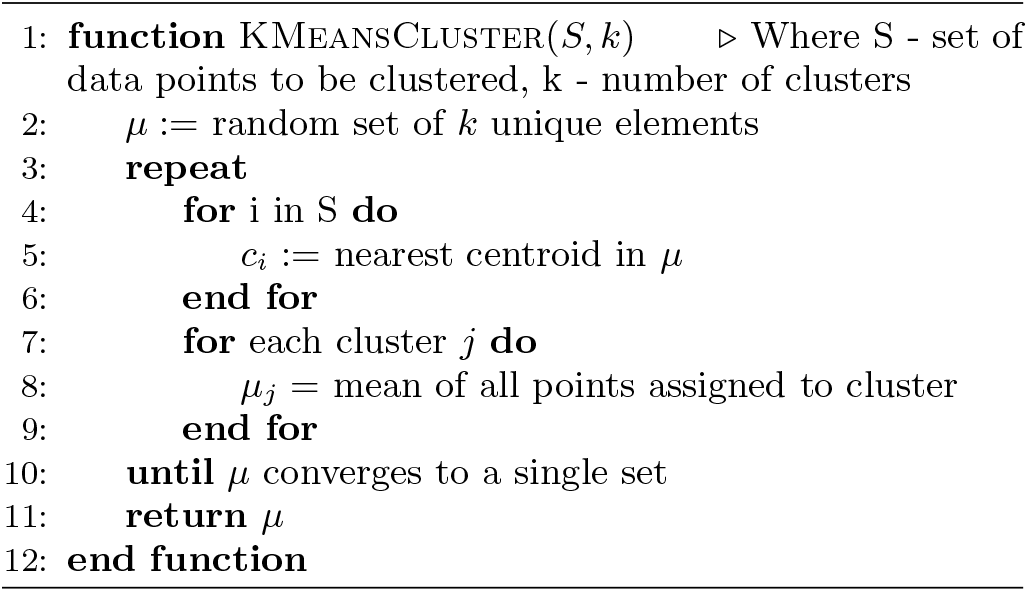

To surmount this problem, we begin our K-Means clustering scheme with a non-randomly chosen set of OPs to act as the vector of initial centroids. These centroids are chosen as to maximize the dissimilarity between the selected OPs. We achieve this by constructing a dissimilarity matrix *A* of *k* OPs that tracks the distance between every pair of OPs within its set. This procedure is summarized in Algorithm 3, and we now describe it in detail.

#### Algorithm 2 K-Means Clustering (Restricted Centroids)

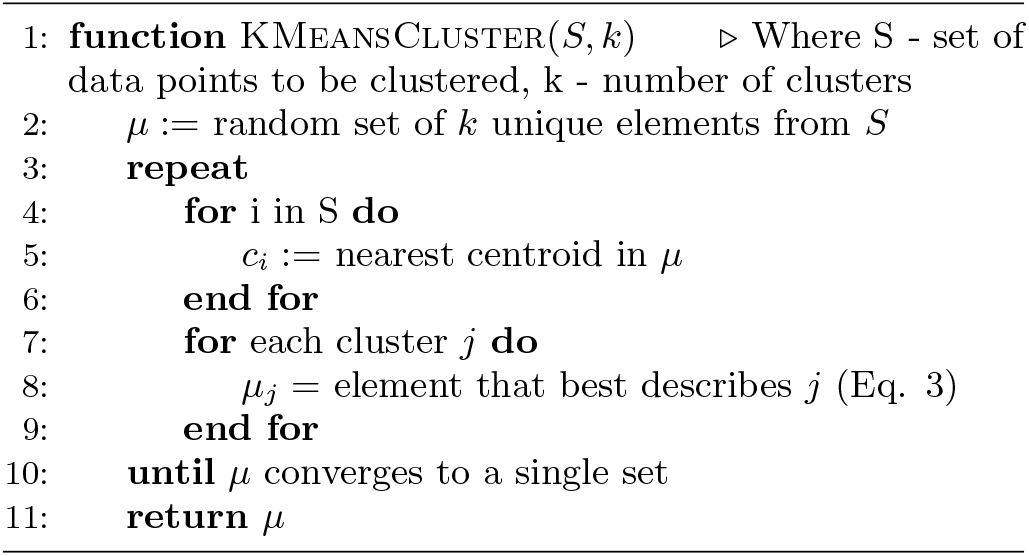

For any dissimilarity matrix *A* of size *k*, which is defined for a given set of *k* OPs, *x*_1_, …, *x*_*k*_, we set any element in the matrix *A*_*i,j*_ ≡ *D*(*x*_*i*_, *x*_*j*_). As a result of this definition a few noteworthy properties arise:

1. *A* is a hollow matrix with all diagonal terms 0, since *D*(*x*_*i*_, *x*_*i*_) = 0 *∀i*.
2. *A* is a symmetric matrix, since *D*(*x*_*i*_, *x*_*j*_) = *D*(*x*_*j*_, *x*_*i*_) *∀i*.
3. The geometric mean of all non-diagonal elements of a row corresponding to an OP gives a measure of how “dissimilar” the OP is to the rest of the set of OPs described by the dissimilarity matrix.

To further elaborate on the third point above, consider an OP *i* and a set *S* of OPs containing *i* and other OPs. The geometric mean *t*(*i, S*) defined in Eq. 4 is a measure of how dissimilar an OP *i* is from the rest of the OPs in the set *S*:

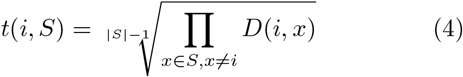

We choose the geometric mean over the arithmetic mean so that OPs that are very similar to other OPs as per the dissimilarity matrix are strongly favored against. As an example, consider the case where two identical OPs are in the full set of OPs. These two OPs would have a distance of 0, yet if the arithmetic mean were to be used, both of these OPs may be included in the dissimilarity matrix if they are very different from the other dissimilarity matrix OPs. However, the geometric mean of the rows corresponding to these OPs would be 0 as long as they are both included, so this procedure would strongly favor removing one of them, which is exactly what we seek.

When considering whether an OP should be added to the dissimilarity matrix, *t*(*i, S*) from Eq. 4 can be used to determine whether the candidate OP should replace an existing OP. Refer to the pseudocode in Algorithm 3 for a description of how the dissimilarity matrix is constructed. It should be noted that this approach is a so-called greedy algorithm^14^ and is not guaranteed to find the *k* most dissimilar OPs. However, this approach does generate a set of OPs that is extremely likely to have higher internal dissimilarity than a randomly chosen set of the provided OPs. Thus, these OPs serve as a good starting point for clustering in the constrained centroid context.

To reiterate, we begin with a very large set *S* of OPs. To determine the *k* best OPs to describe *S* for some *k*, we start by constructing a dissimilarity matrix of size *k* for the set *S* (Algorithm 3). We then use the resulting *k* OPs as the starting centroids for restricted centroid K-Means clustering (Algorithm 2), and the final set of OPs that the solution converges to is the final set of OPs for the given *k*. The approach described up to this point relies on a user input for *k*, the number of output OPs. However, the procedure we have outlined thus far makes no claims about what the best *k* is for a given set *S*. To estimate the optimal *k*, which is the final aspect of AMINO, we turn to rate distortion theory.^10,13^

### C. Jump Method for Determining Number of Clusters

Sugar and James introduced a method for finding the ideal number of clusters for a given dataset using rate distortion theory.^10,13^ An outline pseudocode of their approach is provided in Algorithm 4. The basis for their approach is that once a distortion function is constructed for a data set (a function that captures the error in using some subset of the full dictionary of order parameters to describe the full set), the maximal jump in distortion alone is not the best factor in determining the ideal number of clusters. However, with a proper selection of negative exponent 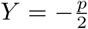 (where *p* is the dimensionality of the system) applied to the distortion function, the largest jump in such an exponentiated distortion corresponds to the optimal number of clusters *k* for the data set. We now explain and quantify this intuitive idea.

In order to apply this approach, a distortion function for a set of centers describing a dataset must be defined. In Euclidean spaces, the Root-Mean-Square-Deviation (RMSD) of distances is typically used.^15^ In a similar fashion using a RMSD built on the distance metric for OPs we just defined (Sec. II A), a distortion function *d*_*k*_ for a set of OPs *S* with centroids *c*_1_, *c*_2_, …, *c*_*k*_ can be written as follows:

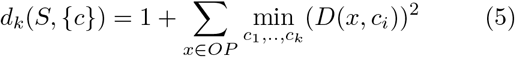

#### Algorithm 3 Dissimilarity Matrix Construction

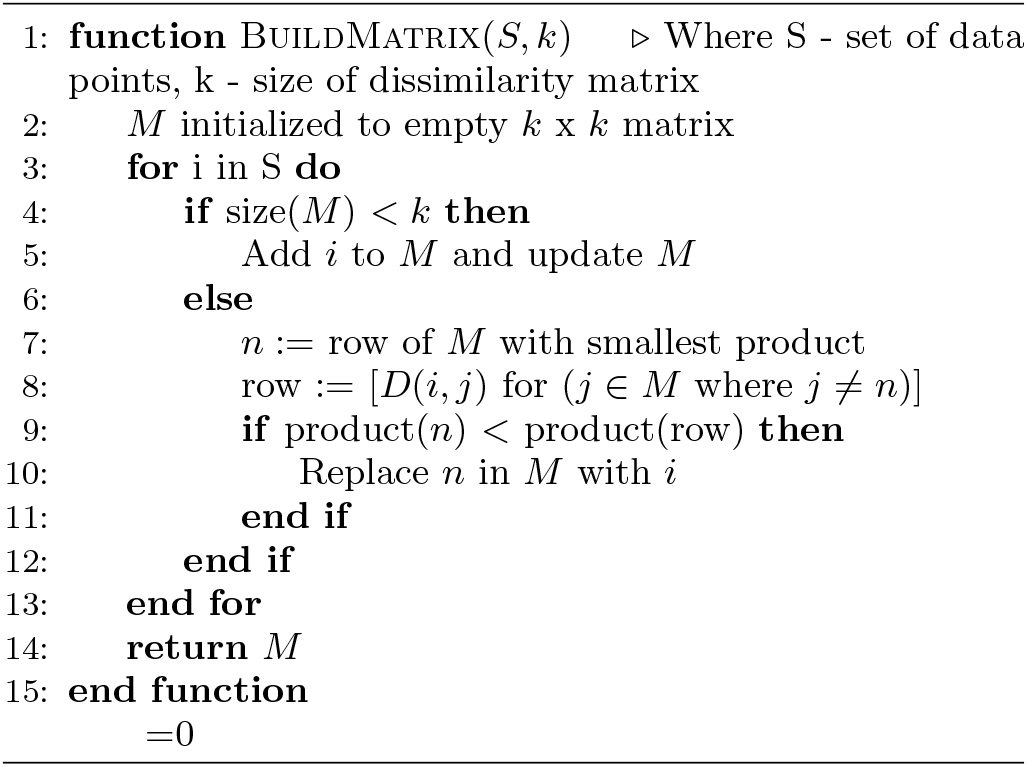

For each value of *k* being tested, *k* OPs are selected using the clustering procedure detailed in Sec. II B. Then, the distortion is calculated as specified in Eq. 5. This distortion represents how well the *k* OPs characterize the entire set of OPs. Then, using Eq. 6, we find the value of *k* that maximizes the jump in the distortion with a negative exponent applied:

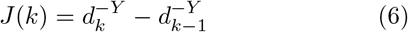

with

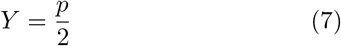

where in Eq. 7, *p* is the dimensionality of the data. However, as stated earlier, OPs lie in an arbitrary higher-dimensional space, and all we have defined so far is a distance function between OPs, not a low-dimensional coordinate system to uniquely identify OPs. Here we would like to highlight that the original work of Ref. 13 recommends setting *p* as the number of independent components contributing to the data. For a protein -ligand system in explicit water comprising, we can say with confidence that 1 < *p* ≪ *N*. That is however not a very useful statement and leaves us guessing regarding the choice of *p*. Indeed recent work has suggested that the degrees of freedom for protein systems is likely fewer than 5.^16–18^ In AMINO, after applying negative exponents corresponding to *p* = 1, 2, .., in 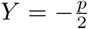, a conservative selection is made by choosing the largest *k* for which a maximal jump is seen in any of these dimensions. Thus, by testing multiple values of *Y*, a value for *k*, the number of clusters, is chosen. Thankfully, as will be shown in the Results section, most selections of *Y* yield fairly consistent results for the best *k*, demonstrating the robustness of AMINO with respect to the precise choice of *p*.

## III. RESULTS

We now provide a series of examples demonstrating clearly how AMINO works as well as its potential usefulness. The examples are systematically constructed, including analytical model potentials and computing binding free energy for a drug fragment and FKBP protein. In the analytical systems we know the true “best” set of OPs beforehand, as they were used to generate the data. To these we then add a large number of decoy OPs and then use AMINO to see if we recover the true OPs. For the protein-ligand system, we consider all possible protein-ligand carbon atom contacts, and then apply AMINO. We use the OPs from AMINO as an input in the reaction coordinate optimization method SGOOP.^4,5^ The RC from SGOOP is then used in metadynamics simulations for binding free energy estimation.

### Algorithm 4 Jump Method to Find k

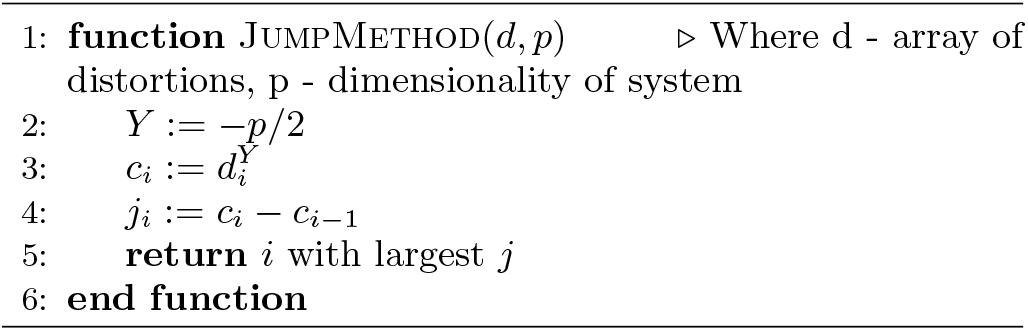

### A. Five Order Parameter Analytical Model System

#### 1. Description

The first analytical model system begins with a collection of 48 OPs, including true OPs and decoy OPs. Each of the 48 OPs fall in one of the following categories:

1. 5 independently sampled random values *A, B, C, D*, and *E* from different distributions with varying numbers of local maxima, all spanning the range [−1, 1]. These are our true OPs and everything else here onward for this system is a decoy OP.
2. 30 linear combinations (including scalar multiples) of these independently-sampled values.
3. 24 versions of one of the random values with random noise added. For clarity, take the example “noisy” OP *A*_1_ ≡ *A* + *q*_1_, where *q*_1_ is a randomly sampled value in the range [−0.05, 0.05]. Each value *A, B, C, D*, and *E* has 3 to 5 different noisy versions of themselves within the set of 48 OPs, and the noise *q*_*i*_ of each noisy version is independent from the noise *q*_*j*_ of any other noisy version.

The end goal is that AMINO should be able to return the original 5 values *A, B, C, D*, and *E* or their scalar multiples. However, noisy versions should not be returned. Although *A* and *A*_*i*_ give the same amount of information about each other, *A*_*i*_ gives less information about *A*_*j*_ than *A* does (for some *i* ≠ *j*), since both *A*_*i*_ and *A*_*j*_ were derived from *A*, but their added noises are uncorrelated. This is not true for scalar multiples, since knowing any scalar multiple of an OP is equivalent to knowing the original OP itself. To help develop intuition for this system, a few distance values are provided in Table I. It can be seen here that *A* has 0 distance from *A* and from −2*A*, but is increasingly more distant from its noisy version *A*_3_ and from the completely different OP *B*.

**TABLE I:** Typical distance values between some of the OPs in the first analytical model system

| OP 1 | OP2 | MI-Based Distance |
| --- | --- | --- |
| $A$ | $A$ | 0 |
| $A$ | $-2A$ | 0 |
| $A$ | $A_3$ | 0.3293 |
| $A$ | $B$ | 0.9957 |

#### 2. Dissimilarity Matrix

It is important to note that dissimilarity matrices were computed for different numbers of clusters, denoted *k*, in the range [1, 30], but only certain dissimilarity matrices of interest are shown. The most interesting cases are for *k* = 4, 5, 6, because these are the values of *k* in a close range of the true value of *k*, which is *k* = 5. The results of the dissimilarity matrix construction (Algorithm 3) are shown in Table II. These OPs are not the final OPs for each *k*, rather, they are the starting points for the restricted centroid K-Means clustering that will be run next. For *k* = 4, the distribution of *E* is not captured by the selected OPs, whereas for *k* = 6, a linear combination of three original OPs was selected by Algorithm 3 as the extra cluster.

**TABLE II:** Result of dissimilarity matrix construction (Algorithm 3) for *k* = 4, 5, 6

| Number of clusters $k$ | OPs resulting from Algorithm 3 |
| --- | --- |
| 4 | $A_1, B_2, C_5, D_5$ |
| 5 | $A_1, B_1, C_3, D_5, E_5$ |
| 6 | $A_3, B_1, C_5, D_3, E_8, A + 4B + E$ |

#### 3. Clustering

For each value of *k*, the OPs resulting from dissimilarity matrix construction (Algorithm 3) were used as the starting centroids in restricted centroid K-Means clustering (Algorithm 1). The final solution that Algorithm 1 converged to for *k* = 4, 5, 6 is presented in Table III.

**TABLE III:** Result of clustering (Algorithm 1) using starting point from dissimilarity matrix for *k* = 4, 5, 6

| Number of clusters $k$ | Post-Clustering OPs |
| --- | --- |
| 4 | $2A, -2B, C, 5D$ |
| 5 | $2A, -2B, C, 5D, E$ |
| 6 | $2A, -2B, C, 5D, E, A + 4B + E$ |

As Table III shows, clustering did return the intended OPs. To clarify why returning scalar multiples falls under intended behavior, consider if the scalar multiple 2*A* was in fact the originally sampled value and that all other scalar multiple and noisy versions were based off of this value instead of *A*. It is impossible to distinguish exactly which of these two explanations is correct (without looking at the code used to generate the distributions of course), and thus, this should fall under intended behavior.

#### 4. Jump Method

Now, using the resulting clusters from running these two steps for different number of clusters *k* in the range [1, 30], we will apply the ideas of Sugar and James^13^ (Algorithm 4) to determine what is the best number of clusters, or equivalently, the number of OPs for this system is. From the description of the system, the answer should be 5. As stated earlier, we do not know the dimensionality of the OP space, so multiple exponents (*Y* in Eq. 6) will be applied. The results can be seen in Fig. 2.

**FIG. 2:**
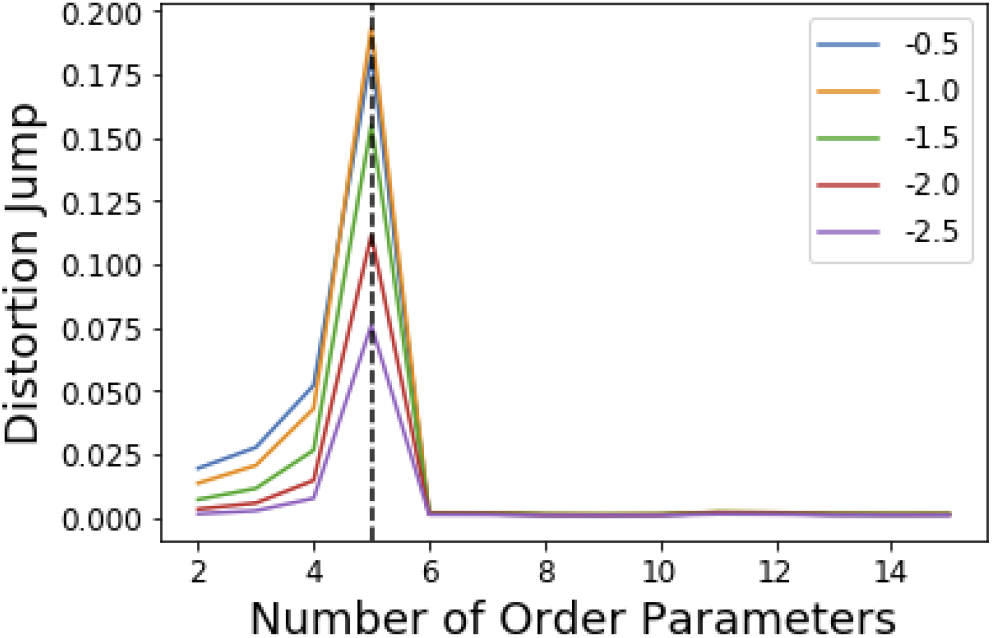
Jumps in distortion with various negative exponents *Y* (indicated with different colors) applied for the 5 OP model system. The black dashed line shows the location for maximum jump at 5 OPs, which is robust with choice of *Y* in Eq. 6.

There is clearly a significant jump at *k* = 5 that corresponds to the correct number of OPs, thus, the *k* = 5 clustering results are used. As stated in Sec. II C, Sugar and James’s results^13^ show that this largest jump in distortion once a negative exponent is applied corresponds to the optimal number of clusters for the data.

### B. Ten Order Parameter Analytical Model System

#### 1. Description

A similar model system as Sec. III A was set up with 10 originally-sampled OPs among a total of 120 OPs. We will refer to the 10 originally sampled values as *A, B, C, …, I, J*, and we will use the same conventions described in Sec. III A to describe the noisy versions of these values.

The OPs in this data set consisted of:

1. 10 independently sampled random values *A, B, C, …, I, J* from different distributions with varying numbers of wells, all spanning the range [*−*1, 1].
2. 110 “noisy” versions of one of the random values with random noise added.

For this system, we did not add any linear combinations and instead focused solely on noisy versions of the original OPs, since these are more similar to what would be encountered in an experimental system. Similar to Sec. III A, we find that AMINO recovers the originally-sampled 10 OPs, since these 10 give the most information about the entire system, or equivalently, the originally-sampled 10 OPs provide the most information about the full set of 120 OPs. Since there are no scalar multiples in this system, there are no pairs of OPs with a distance of 0 (as determined by Eq. 2), unlike in the 5 OP model system from Sec. III A.

#### 2. Dissimilarity Matrix

Dissimilarity matrices were computed for *k* in the range [1,30], and the values provided in Table IV are the results of dissimilarity matrix construction algorithm (Algorithm 3) for the values of *k* near the true number of clusters, *k* = 10.

**TABLE IV:** Result of dissimilarity matrix construction (Algorithm 3) for *k* = 9, 10, 11.

| Number of clusters $k$ | Dissimilarity Matrix OPs |
| --- | --- |
| 9 | $A_{13}, B_1, C_4, D_4, E_{13},$<br>$G_3, H_2, I_{11}, J_1$ |
| 10 | $A_{11}, B_8, C_6, D_4, E_8, F_8,$<br>$G_{12}, H_2, I_9, J_1$ |
| 11 | $A_9, B_1, C_4, D_3, E_{12}, F_5,$<br>$G_8, H_2, I_3, J_1, J_8$ |

#### 3. Clustering

As was done in Sec. III A, the results of Algorithm 3 in Table IV were used as input to Algorithm 2 in order to find the final centroid OPs that AMINO will return. The results of clustering for the interesting values of *k* are in Table V. Just as in Sec. III A, the original OPs are returned for the correct value of *k*, which in this case is *k* = 10, and deviating from the correct value of *k* either leads to missing an original order parameter or selecting a noisy version.

**TABLE V:** Result of clustering (Algorithm 1) using starting points from dissimilarity matrix for *k* = 9, 10, 11

| Number of clusters $k$ | Post-Clustering OPs |
| --- | --- |
| 9 | $A, B, C, D, E, G, H, I, J$ |
| 10 | $A, B, C, D, E, F, G, H, I, J$ |
| 11 | $A, B, C, D, E, F, G, H, I, J, J_3$ |

#### 4. Jump Method

Now, we look at the resulting jumps in distortion from using different values of *k* for the 10 OP model system. Fig. 3 shows that the selected number of OPs as determined by Eq. 6 is *k* = 10, as expected given that there were 10 initial OPs.

**FIG. 3:**
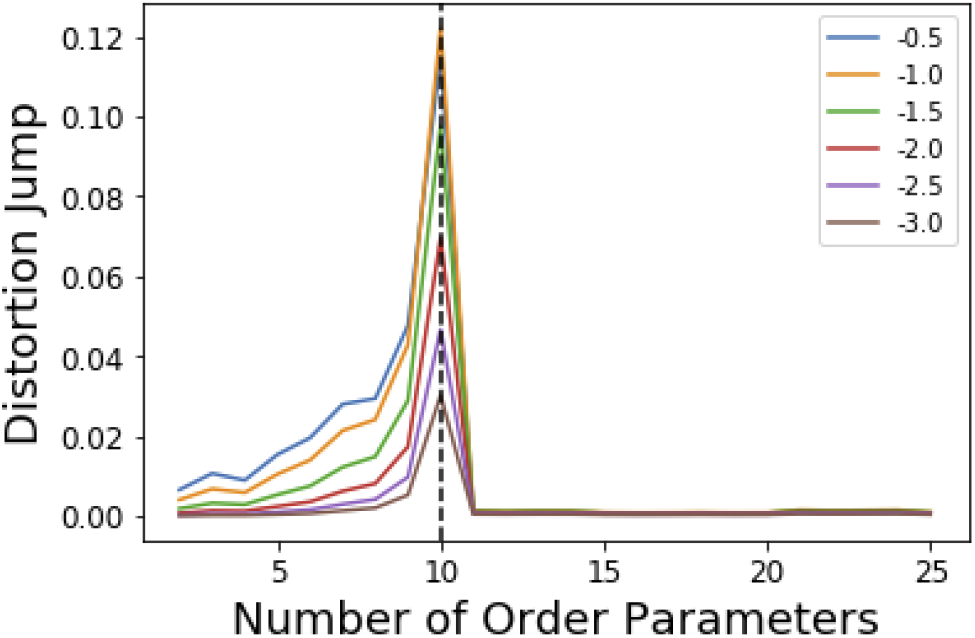
Jumps in distortion with various negative exponents *Y* (indicated with different colors) applied for the 10 OP model system. The black dashed line shows the location for maximum jump at 10 OPs, which is robust with choice of *Y* in Eq. 6.

As shown here in Sections III A and III B, this approach appears robust in handling these kinds of model systems. Now, we consider OPs extracted from unbiased MD trajectories of real systems to determine whether this procedure remains successful.

### C. FKBP/BUT System

Metadynamics^8,19^ is a popular sampling method used to bias MD simulations in order to visit states separated by a large energy barrier that would rarely be traversed in an unbiased MD simulation. The biasing is performed along a pre-selected low-dimensional RC. The typical output of a metadynamics simulation is the equilibrium probability distribution of the system along the RC or along any other low-dimensional coordinate through a reweighting scheme.^19^ There also exist variations of metadynamics useful to construct not just static probabilities but also unbiased kinetic observables such as the rates for moving between different metastable states.^20,21^ In any of its formulation, metadynamics benefits from a suitably chosen RC that adequately characterizes the different states of interest.^8^ While of late many methods have been developed that generate a RC from a set of OPs^3,4,6,7,22^, as stated earlier, the input set of OPs is usually still chosen based on prior knowledge of the system. However, by applying AMINO, this set of OPs can be automatically selected with minimal prior knowledge of the system, except the bound pose structure and a short unbiased MD run where we do not rely on the dissociation event being sampled even once. In order to compute the free energy of binding/unbinding, several dissociation and re-association events need to be observed in order for the free energy to converge. However, using metadynamics with a properly-constructed RC, unbinding events can occur more quickly (in terms of simulation time), decreasing the total time needed to compute the free energy.

We begin with a short 10 nanosecond trajectory of the protein-ligand system of the protein FKBP and one of its ligands, 4-hydroxy-2-butanone (BUT). The trajectory is expressed in terms of a dictionary of 428 OPs (Fig. 4) that consists of every combination of distances between alpha carbons in the protein (107 atoms) and carbons in the ligand (4 atoms). These 428 OPs were used as input to AMINO to yield a reduced dictionary of 8 OPs, shown in Table VI. The output of using the jump method for this system is illustrated in Fig. 5 where irrespective of the precise value of *p* we can identify 8 OPs as the robust choice. Table VI also provides the weights obtained for these OPs when considered in a 1-dimensional RC in SGOOP^4^ expressed as a linear combination of these OPs. Now that a reaction coordinate has been constructed using the AMINO-selected OPs, a biased metadynamics runs was conducted to calculate the free energy of binding (∆G) for the system. We provide the results from the metadynamics run as well as for an unbiased run for comparison in Fig. 6.

**FIG. 4:**
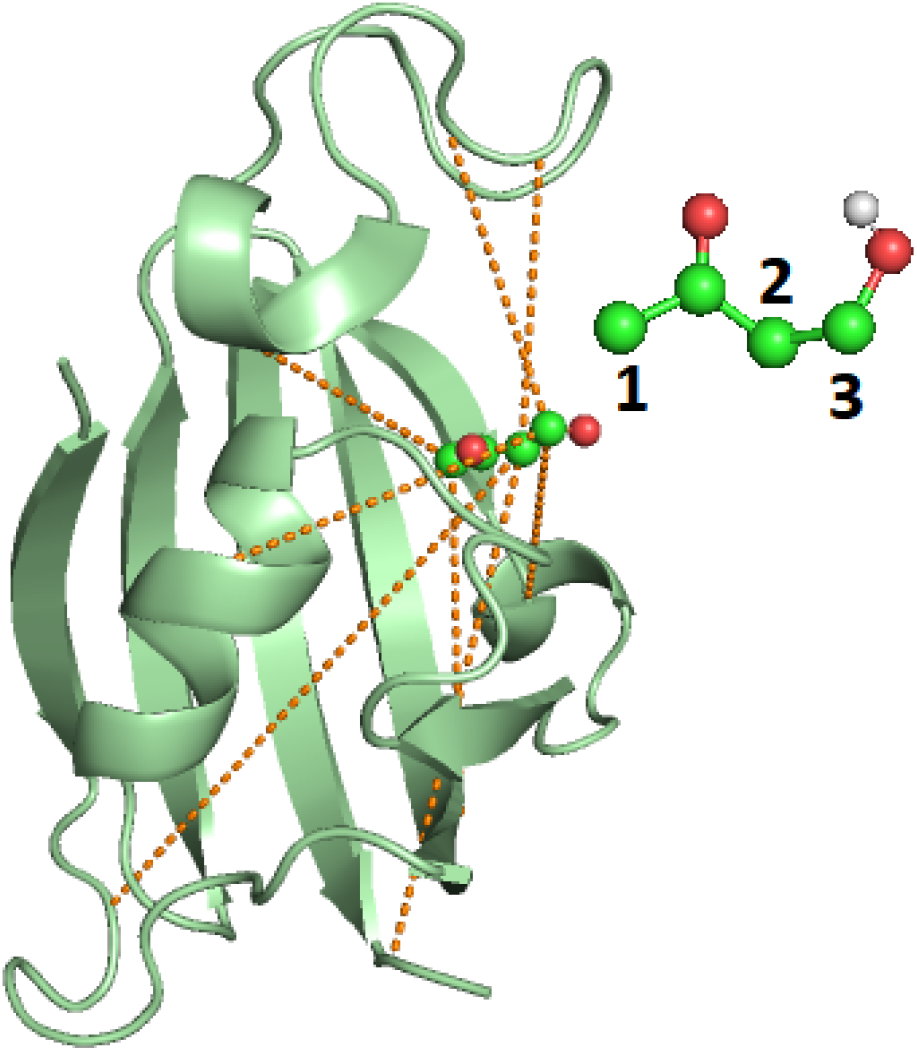
Image of the bound FKBP/BUT protein-ligand system, with inset showing the atom numbering for ligand carbons used in Table VI. Carbons are in blue, oxygens are in red, and the only polar hydrogen is in white. The dashed lines show the 8 different OPs detailed in Table VI, out of input 428 options, resulting from AMINO that were then used to construct a RC through SGOOP^4^

**FIG. 5:**
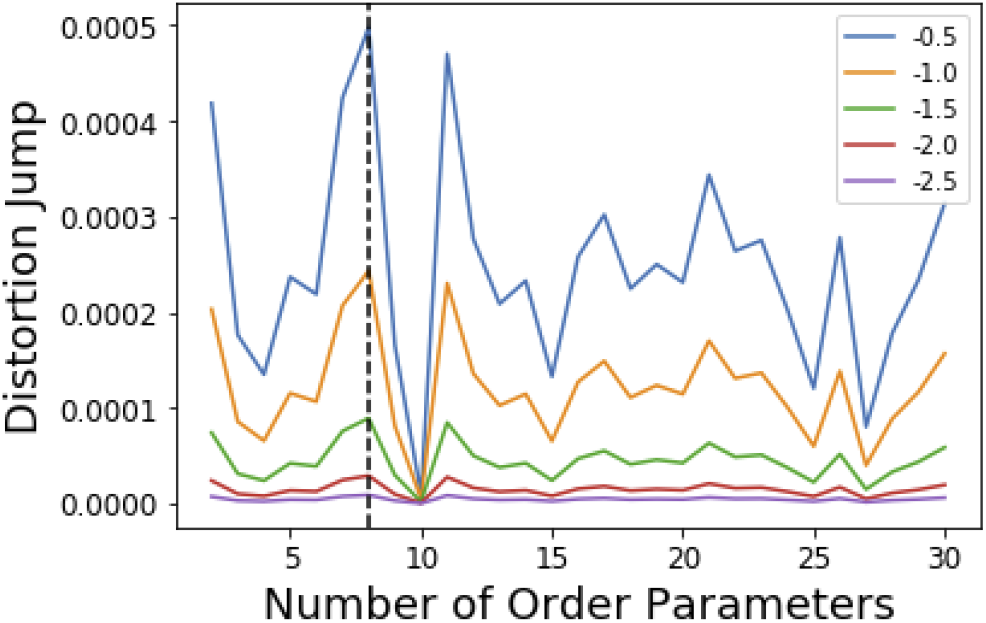
Results of the jump method (Algo 4)) when applied to the FKBP/BUT system. The different lines correspond to different choices for *Y* in Eq. 6, specified in the legend. The maximum jump occurs at 8 OPs for all choices of *Y*.

**FIG. 6:**
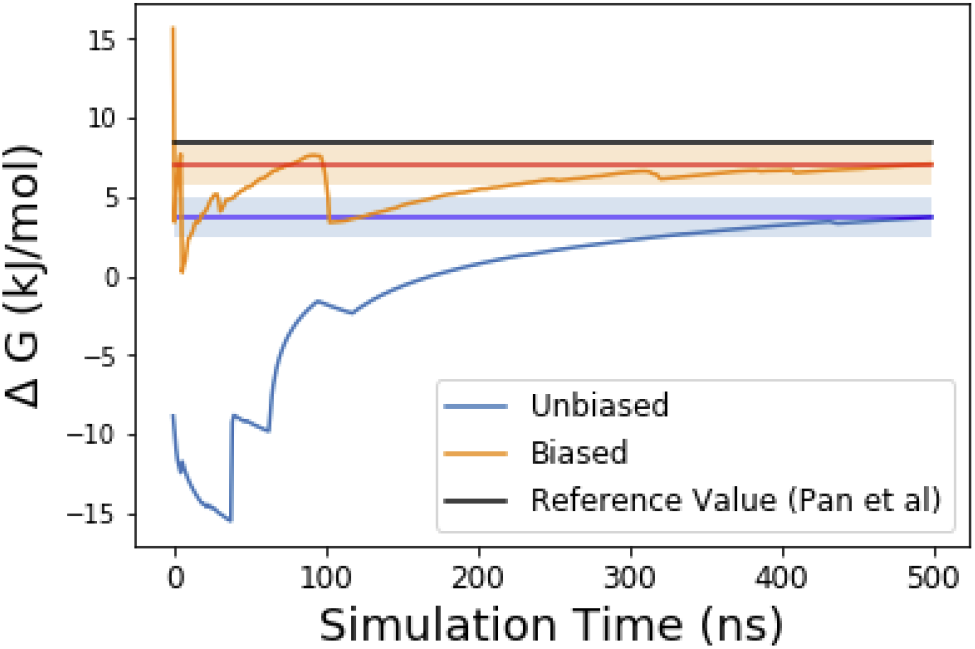
Absolute binding free energy ∆G in *kJ/mol* as a function of simulation time for the unbiased and biased runs of the FKBP/BUT protein-ligand system. The shaded regions denote a ±*k*_*B*_*T*/2 range from the final ∆G estimate. The horizontal solid blue and orange lines denote respectively to the final ∆G values after 500 ns in the unbiased and biased simulations, respectively. Solid black line shows the reference value from long unbiased MD simulations performed on Anton supercomputer by Pan et al.^23^

**TABLE VI:** List of OPs that were output from AMINO for FKBP/BUT protein-ligand system. Ligand carbon numbers are shown in Fig. 4

| Protein Atom | Ligand Atom | Weight in SGOOP RC |
| --- | --- | --- |
| 86GLY Cα | 3 | 1.000 |
| 106LEU Cα | 2 | 0.946 |
| 39SER Cα | 3 | -0.973 |
| 23VAL Cα | 1 | -0.862 |
| 87HIS Cα | 2 | -0.903 |
| 61GLU Cα | 3 | -0.951 |
| 10GLY Cα | 2 | -0.974 |
| 80TYR Cα | 1 | -0.846 |

Fig. 6 shows how the simulation that was biased using a RC constructed from AMINO-generated OPs gives a reasonably accurate estimate of ∆*G* in a very short time. Also illustrated is the reference value for this system from long Anton simulations reported in Ref. 23. The difference between our estimate and the Anton value is minuscule especially considering that we re-parameterized the ligand on our own. While we followed Ref. 23’s instructions, this can easily lead to the fraction of a *k*_*B*_*T* difference we find. On the other hand our unbiased MD simulation gives a much worse estimate of ∆*G* and as can be seen from Fig. 6 as well as Fig. 7, it would take much longer than 500 ns to converge to the correct value. Thus, the OPs selected by AMINO using absolutely no prior knowledge of the system were able to construct a meaningful reaction coordinate that led to enhanced sampling of this system.

**FIG. 7:**
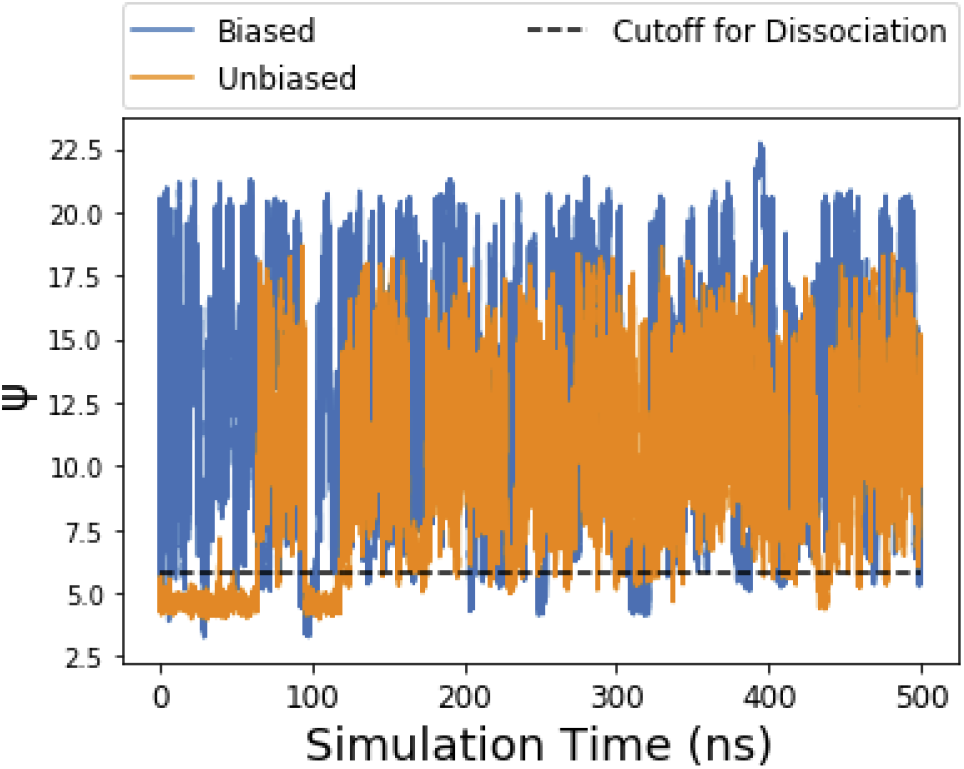
Time-series of reaction coordinate obtained from SGOOP as a linear combination of AMINO-outputted OPs. Blue lines show biased simulations with multiple dissociation and association events leading to quick estimate of converged ∆*G*, while orange lines show unbiased simulation where the trajectory stays trapped for extended periods of time leading to poor estimates of ∆*G*. The black dashed line indicates the boundary between bound and unbound considered by us to stay consistent with the definition of bound state in Ref. 23.

Although metadynamics does significantly increase the occurrence of unbinding events, re-binding is limited by entropy, thus traditional metadynamics^8^ cannot encourage re-binding, regardless of the reaction coordinate. This is illustrated in Fig. 7. The unbiased trajectory begins in the bound state, and the ligand only ever rebinds to the protein once. In the biased run, the simulation spends very little time in the bound state since it is biased away from remaining in the already-explored bound state. This allows the simulation to spend more time searching the other, unexplored states, so that it converges to a reasonable solution more quickly. However, this extended time spent in the unbound state presents an opportunity for further acceleration through approaches such as funnel metadynamics which help with the entropic problem.^24^ In combination with AMINO, SGOOP, and traditional metadynamics, funnel metadynamics can even further accelerate the procedure we have gone through here.

## IV. DISCUSSION

In this work we have introduced an information theoretic approach to screening for OP redundancy by using a mutual information based distance function as a measure of dissimilarity between two OPs. In general, to select a set of OPs for a system, current approaches rely primarily on previous biophysical knowledge of the system. With the procedure we have presented, a set of viable OPs can be constructed with much less knowledge of the system. For the calculation of protein-ligand absolute binding free energy, this amounted to knowing the bound pose structure and a very short MD trajectory (10 ns) where the ligand did not have to dissociate even once. Having a trajectory with actual dissociation events would likely increase the accuracy of AMINO even further, but that is not a practical scenario for systems of real-world interest invariably plagued with the sampling issue. The approach applied involves clustering the set of OPs by using the mutual information based metric as detailed in Algorithm 2, and using the resulting centroids as the output set of OPs. In order to overcome the problem of centroids becoming stuck in the same cluster, we initialize the K-Means clustering algorithm with centroids from the construction of a dissimilarity matrix (Algorithm 3). The motivation for creating the dissimilarity matrix is to generate a set of *k* points out of the set of provided OPs that are internally dissimilar. To determine the best *k*, a rate distortion theory based Jump Method, described by Sugar and James,^13^ was employed on the results of clustering for various *k*.

Our proposed algorithm eases the previous requirement of how well a system must be understood before selecting OPs. Selection of OPs is vital to all methods employed after unbiased MD simulations, such as reaction coordinate construction and enhanced sampling. In future work, we will be applying AMINO to different molecular systems in biology and beyond, with different classes of trial OPs, including bond torsion values, hydration states of specific constituents and many others. We believe that our algorithm provides significant progress towards the process of automating OP selection. A Python 3 implementation of this algorithm in Jupyter Notebook is available at https://github.com/pavanravindra/amino for public use.

## ACKNOWLEDGEMENTS

We thank Deepthought2, MARCC and XSEDE (projects CHE180007P and CHE180027P) for computational resources used in this work. We would also like to thank Shashank Pant for help with setting up the FKBP/BUT system for MD simulation.

